# Group social conditions and environment predict foraging behavior in wild baboons

**DOI:** 10.1101/2025.11.20.689548

**Authors:** Maria J.A. Creighton, Olivia Fan, J. Kinyua Warutere, Jenny Tung, Elizabeth A. Archie, Susan C. Alberts

**Affiliations:** Department of Biology, Duke University, Durham, North Carolina, USA; Amboseli Baboon Research Project, PO Box 72211-0020, Nairobi, Kenya; Department of Evolutionary Anthropology, Duke University, Durham, North Carolina, USA; Duke University Population Research Institute, Duke University, Durham, North Carolina, USA; Department of Primate Behavior and Evolution, Max Planck Institute for Evolutionary Anthropology, Leipzig, Germany; Faculty of Life Sciences, Leipzig University, Leipzig, Germany; Department of Biological Sciences, University of Notre Dame, Notre Dame, Indiana, USA

## Abstract

For group-living animals, conditions in the physical and social environments are closely linked to foraging outcomes, but the nature and causal direction of many aspects of these relationships remain unclear. Here, we use long-term data from a well-studied population of wild baboons in Amboseli, Kenya, to examine how group-level social traits (group size and social network density) and climate variables (rainfall and temperature) are linked to two types of foraging outcomes for adult female baboons: (i) foraging-related time budgets and (ii) diet composition (time spent on fallback foods, such as grass corms, versus high-energy foods). We find that rainfall and temperature interact to predict multiple foraging outcomes: more rainfall is associated with more time spent feeding, less time spent walking without feeding, and more feeding time spent on grass corms, but this influence is more pronounced in hotter years than in cooler years. Females in intermediate-sized groups spend more time walking without feeding than those in other groups, but group size does not significantly predict other foraging-related outcomes (i.e., time spent feeding or diet composition). We also find that females in groups with denser social networks spend less time feeding, and less time feeding on grass corms, than those in sparser networks. However, in a preliminary causal analysis meant to further explore this result, we find support for the hypothesis that this relationship is driven by effects of foraging on social behavior, as opposed to effects of network density on foraging: more time spent eating grass corms leads to lower social network density and more time spent feeding. Our results show how the social and the physical environments are linked to foraging outcomes, and highlight the importance of future studies of their pattern and mechanism.

## INTRODUCTION

Variation in foraging behavior can have a substantial impact on fitness. For instance, individual variation in foraging behavior predicts survival and/or reproduction in species as diverse as Darwin’s finches (Grant 1985), juvenile redshanks (Sansom et al. 2009), water striders (Blanckenhorn 1991), wandering albatrosses (Patrick & Weimerskirch 2015), Soay sheep (Illius et al. 1995), and infant baboons (Altmann 1991). Given these effects on fitness, understanding the forces that explain individual variation in foraging outcomes is an important topic in animal ecology. Conditions in the physical and social environment are both thought to be important in shaping foraging behavior but additonal work is required to fully understand the magnitude and direction of these relationships.

With respect to environmental conditions, individuals in many species adjust aspects of foraging effort — such as travel distances, activity budgets, and diet composition — in response to changes in rainfall and/or temperature, which frequently determine food availability. Given that both temperature and rainfall vary hugely across ecosystems, expected effects of these variables on foraging-related behaviors depend on the study system in question. Generally, rainfall promotes the growth of important food items for many terrestrial species, resulting in species-specific effects on foraging behavior. For instance, the effects of rainfall on foraging-related behaviors can vary by age and sex (gorillas, Auger et al. 2023), by food type (brown lemurs, Sato et al. 2014; geladas, Fashing et al. 2014), and by social group (baboons, Bronikowski & Altmann 1996; Chowdhury et al. 2021). Meanwhile, cooler temperatures are often associated with increased time invested in feeding (Barbary macaques, Majolo et al. 2013; vervet monkeys, McFarland et al. 2014; bamboo lemurs, Eppley et al. 2016; baleen whales, Owen et al. 2018) and increased foraging capacity (banded mongooses, Khera et al. 2025). In climates where high temperatures exceed an animal’s thermal neutral zone, individuals might decrease time spent feeding in favour of more sedentary behaviors like resting to prioritize thermoregulation (Hill 2006).

Group-level social conditions like group size and within-group social relationships are also linked to foraging behavior. For instance, data from multiple animal species support the idea that group size drives foraging outcomes through multiple processes, including the demands of cooperative foraging (e.g., wild dogs, Creel & Creel 1995; African lions, Packer et al. 1990; killer whales, Baird & Dill 1996), the consequences of group size for within- and between-group competition (e.g., baboons, Hamilton et al. 1975; lions, McComb et al. 1994; capuchin monkeys, Crofoot et al. 2008; reviewed in Markham & Gesquiere 2017), and the time individuals spend being vigilant instead of foraging (e.g., starlings, Powell 1974; ostriches, Bertram 1980; juncos, Caraco 1979; waders, Barbosa 2002). As a consequence of these and other processes, group size is an important predictor of foraging-related behaviors (daily travel distances and time budgets; see Sterck et al. 1997 and Majolo et al. 2008 for meta-analyses in primates). Social relationships within groups have a more complex relationship to foraging than group size. On one hand, flexible group-level social traits such as social network density may shape foraging outcomes if numerous, strong social connections enhance individual foraging ability (hereafter referred to as the “social benefits hypothesis”). At the same time, if animals are required to spend more time foraging, they will have less time to develop and maintain social relationships, creating a trade-off between time spent foraging and time spent socializing (hereafter referred to as the “foraging trade-off hypothesis”). These processes — social benefits enhancing foraging and foraging demands trading off against social time — need not be mutually exclusive.

Evidence from several species supports the social benefits hypothesis. Groups that are socially well-connected for their size may benefit in intergroup competition by cooperating more effectively in intergroup contests, as demonstrated in wild chimpanzees (Samuni et al. 2021). Members of well-connected groups may also experience more opportunities for social learning through increased exposure to conspecifics. For instance, in a meta-analysis of 33 species in five taxonomic groups (invertebrates, fish, reptiles, birds and mammals), individuals were more likely to learn new foraging behaviors from close social partners (Camacho-Alpízar & Guillette 2023). A related result emerges from an experimental study of wild baboons, which demonstrated that the tendency for individuals to follow a “leader” to a food source was mediated by social ties to that leader (King et al. 2011). Similarly, in an observational study of brown lemurs, the joining and following patterns exhibited by group members during foraging were better predicted by affiliative relationships between a leader and a follower than by several other potential mechanisms, including kinship or the number of other individuals following the leader (baboons, King et al. 2011; brown lemurs, Jacobs et al. 2011).

Research also provides support for the foraging trade-off hypothesis. Groups that forage most efficiently or maintain access to high-quality resources may have more time, energy, or social tolerance to invest in affiliative social behaviors (Dunbar et al. 2009). Trade-offs between foraging needs and social time have been demonstrated in multiple species, including red deer (Clutton-Brock et al. 1982), baboons (Altmann 1980), ravens (Pfuhl et al. 2014), and meerkats (Clutton-Brock et al. 1999). Furthermore, a meta-analysis of multiple species of nonhuman primates suggests that animals in large groups are forced to reduce social time in order to meet ecological demands (Lehmann et al. 2007). In other words, variation in food availability can impose constraints on social time budgets, with the result that social relationships within a group will vary with foraging demands.

Using longitudinal data from a well-studied population of wild baboons (*Papio cynocephalus* x *P. anubis*) in the Amboseli ecosystem of Kenya, we investigated the effects of group-level social traits (group size and social network density) and climate variables thought to influence food availability (temperature and rainfall) on annual group-level foraging outcomes. Our first group-level social trait, group size, captures the number of individuals physically present in a group and has been shown to have multiple and conflicting links to foraging-related outcomes in primates. For example, in our study population intermediate sized groups have smaller home range sizes, shorter daily travel distances, lower glucocorticoid concentrations, and more even patterns of space use than the largest and smallest groups (Markham et al. 2015). In contrast, large mountain gorilla (*Gorilla beringei beringei*) groups have higher energy intake despite increased travel (Grueter et al. 2018). Meanwhile, a meta-analysis of primates showed that species with larger groups tend to travel and feed more (Majolo et al. 2008). In addition to group size, network density (i.e., the proportion of dyads in a network that are observed to interact) is another important group-level social trait which reflects how socially connected individuals in a group are to one another. While smaller groups tend to have higher network densities, group size explained a relatively small proportion of the variance in social network density in an analysis of multiple social species (Shizuka & McDonald 2015), indicating that this relationship is not deterministic. Unlike group size, the relationship between network density and foraging has not been empirically explored.

We first tested whether our climatic and social variables of interest predicted foraging outcomes using generalized estimating equations (GEE). Next, in a post-hoc analysis based on the results of our GEE models with respect to social network density, we conducted a causal path analysis to evaluate the relative support for either the social benefits hypothesis or the foraging trade-offs hypothesis in explaining our results. Specifically, our GEEs identified relationships between network density and two key foraging outcomes: time spent feeding and the proportion of feeding time devoted to grass corms (a slow-to-harvest “fallback” food for baboons; Altmann 2009). We therefore considered a simplied set of potential causal relationships between social network density and these two foraging outcomes, as depicted in Figure 1. This simplified schematic does not account for many other variables that affect baboon behavior, but it represents a useful conceptual framework that we use to consider the possible relationships among these three variables.

**Figure 1:**
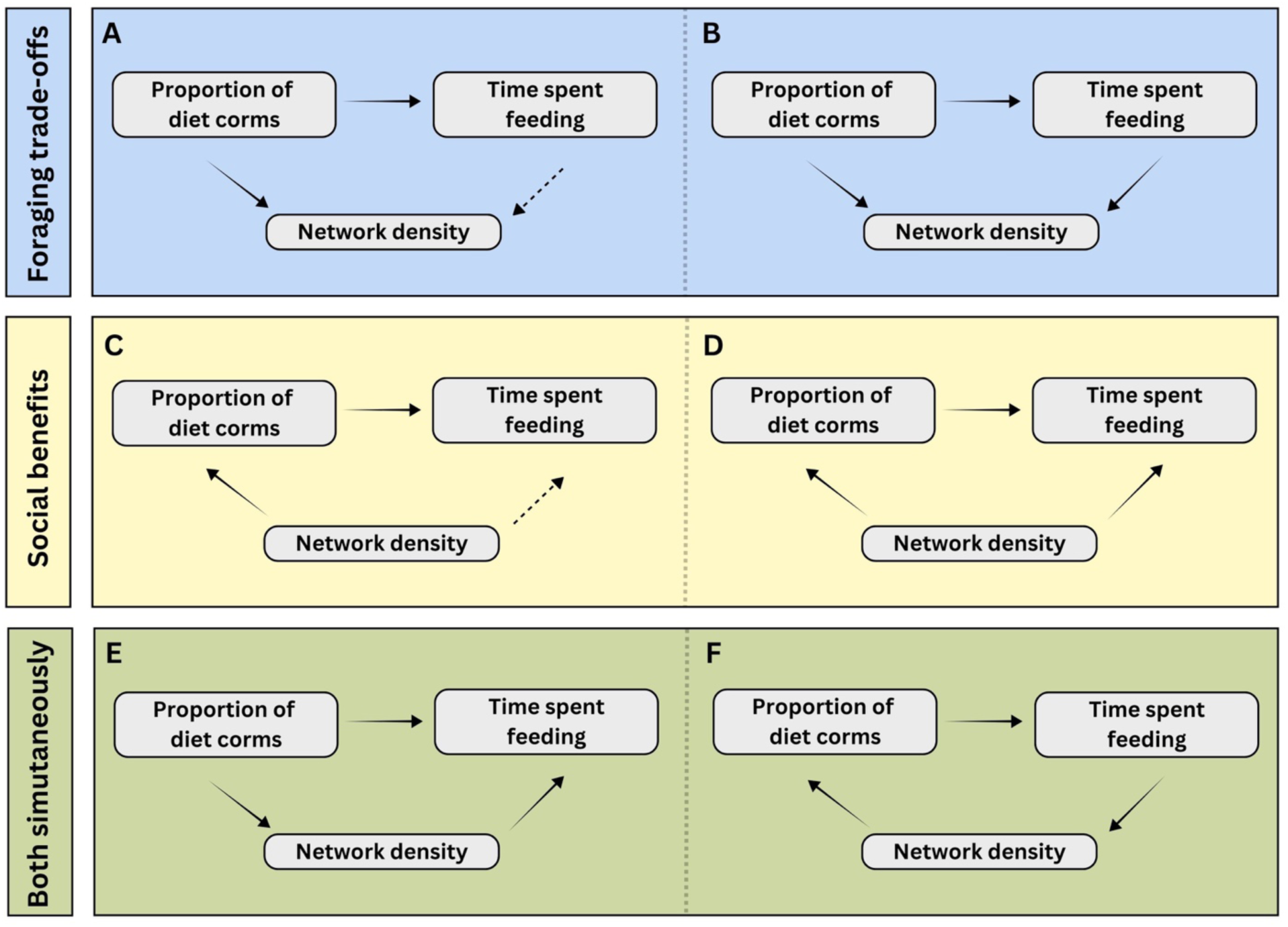
Six possible causal models of the relationship between network density and foraging outcomes; two models are depicted in each panel. In all three panels, environmental variables (temperature and rainfall) affect diet composition (proportion of the diet composed of corms), which in turn is a predictor of increased time spent feeding in this population (hence the arrow pointing from the proportion of corms in the diet to time spent feeding appears in every box). The blue box that encompasses A) and B) depicts possible predicted relationships under the foraging trade-off hypothesis: A) diet composition affects network density creating an indirect correlation between network density and time spent feeding or B) time spent feeding and diet composition both directly affect network density. The yellow box that encompasses C) and D) depicts possible predicted relationships under the social benefits hypothesis: C) network density affects diet composition creating an indirect correlation between network density and time spent feeding or D) nework density directly affects both time spent feeding and diet composition. The green box that encompasses E) and F) depicts a scenario in which both directions of causality are supported: E) diet composition affects network density but network density affects time spent feeding or F) time spent feeding affects network density but network density affects diet composition. Solid arrows represent direct effects whereas dashed arrows represent indirect correlations.

## METHODS

### DATA

#### Study population and study subjects

Wild baboons in the Amboseli basin have been under continuous observation since 1971 as part of the Amboseli Baboon Research Project (Alberts & Altmann 2012; Alberts et al. 2020). The Amboseli ecosystem is a semi-arid savannah environment with the typical annual wet/dry seasonal dynamics of east Africa, which includes two dry and two wet seasons each year (Nicholson 1996). Baboons in this population are admixed with majority yellow baboon ancestry (*Papio cynocephalus*) and minority anubis baboon ancestry (*P. anubis*; also known as olive baboons) (Alberts & Altmann 2001; Tung et al. 2008; Vilgalys & Fogel et al. 2022). The Amboseli Baboon Research Project monitors between two and six social groups (“study groups”) at a time, all of which originated from two original study groups that have undergone multiple fissions. Each study group is visited two to four times a week by experienced observers who are able to visually recognize individual animals. During group visits observers conduct a census where they record all individuals present in the group on that day, and collect behavioral data via focal sampling and representative interaction sampling (described below).

#### Level of analysis

In our investigation of the relationship between group size, network density, climate variables, and foraging behavior, the level of observation was group-years (i.e., each row of the dataset represented one year of a group’s existence). Each group-year started on November 1^st^, the beginning of the short rainy season, and ended on October 31^st^ of the following calendar year, the end of the long dry season. Our dataset included 100 group-years across 13 distinct social groups between 2000 and 2021. We did not consider data from before 2000 because differences in focal data collection protocols before and after 2000 meant that some types of data used for our analysis were not available prior to 2000.

#### Group size

We measured group size as the total number of unique adult individuals in a social group during a given year. In measuring group size, we only counted adults who were present in the group for at least half of the year (183 days). This approach avoided including individuals who were present for a relatively small part of the year and thus did not contribute substantially to group-level dynamics for most of the year (e.g., dispersing males or individuals who died during the group-year). We additionally measured total group size (including juveniles and infants) but this metric led to similar results as adult group size in all analyses. For simplicity we therefore report analyses and results using adult group size throughout.

#### Network density

Social network density (i.e., the proportion of dyadic pairs or “edges” in a network that are observed to interact) was calculated from networks of grooming between adult group members in each group in each year. We considered two individuals to be connected in the network if they were observed to groom at least once over the course of the group-year. Grooming data for this population are collected continuously by observers via “representative interaction sampling” during visits to a group. This protocol is described in Alberts et al. (2020). Briefly, observers record all observations of grooming for any individuals within their line of sight, while simultaneously conducting focal animal follows of adult females and juveniles in a predetermined randomized order. As observers move from one focal individual to another this approach ensures representative sampling of the whole social group (Alberts et al. 2020). To be included in a network, adults needed to be present for at least 183 days of the given year (whether or not they were ever observed to groom). Figure 2A shows an example of an observed network in our study population with low network density and Figure 2B shows an example of an observed network of similar size with high network density. Because each group changed in size and in membership substantially over the years in which they existed, groups exhibited considerable variation in size and density over the course of their existence (Figures S1 and S2). Group size and network density were negatively correlated with one another, but not strongly so (r=-0.386; Figure S3).

**Figure 2:**
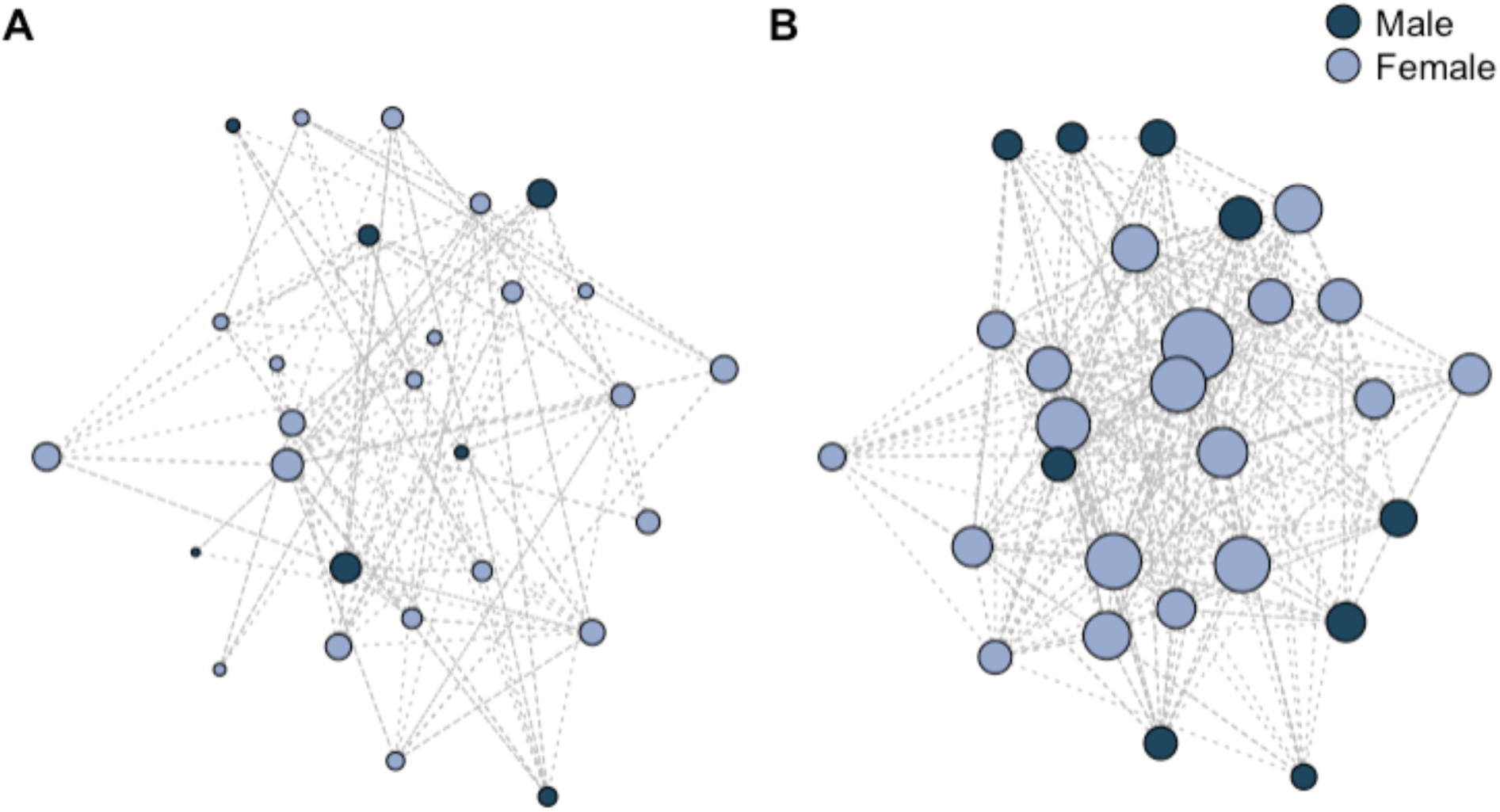
Example of observed networks with comparable adult group sizes but differing network densities. A) Depicts the grooming network of group 1.11 during the 2011 hydrological year (from November 1^st^ to October 31^st^ of the following calendar year). This is an example of a group with a low network density of 0.178. Adult group size for this group was 31, comprising 21 females and 10 males. B) Depicts the grooming network of group 2.20 during the 2003 hydrological year. This is an example of a group with a high network density of 0.521. Adult group size for this group was 28, comprising 19 females and 9 males. In both A and B the size of nodes increases with the number of observed grooming events an individual engaged in and dotted lines connect dyads who have grooming relationships. Note that these examples represent raw network densities (i.e., not corrected for observer effort) and that individuals without any grooming relationships are not pictured (but were included in network density calculations).

#### Environmental data

Rainfall has a large influence on the resources available at any given time of year in Amboseli as it modulates the growth of many food items (Gesquiere et al. 2024). Average daily rainfall in each year (i.e., the average amount of rainfall in millimeters that fell per day within the year of interest) was estimated from daily rainfall measurements that were collected using a rain gauge near the study area (Alberts et al. 2020). Because temperature also has an effect on the physiology and behavior of baboons (Hill 2006; Gesquiere et al. 2008; Gesquiere et al. 2024) and may therefore also influence foraging, we additionally calculated the average maximum daily temperature in Celsius for each group-year using data from a min-max thermometer located near the study area that is checked once every 24 hours.

#### Foraging data

To obtain information about adult female foraging activity, we used observations from 10-minute focal animal samples collected on adult female baboons during visits to each study group. During each focal sample, the subject’s activity (feeding, resting, walking, grooming, being groomed, or other) was recorded every minute as a ‘point’ sample, yielding 10 point samples per complete focal sample (Alberts et al. 2020). When the activity was feeding, point samples also included information about the category of food on which the focal animal was feeding (e.g., leaves, grass blades, corms, fruits, gum, seeds, etc.). We pooled all point samples for all adult females in a social group in a given hydrological year together. Groups had between 1072 and 8315 female focal point samples per year (mean = 4966±1571). Using these pooled point samples from adult female focals, we generated counts of the following for each group-year: (i) the number of total focal points collected that year, (ii) the number of points spent feeding (i.e., when an animal is making contact with, reaching for, or processing food; Alberts et al. 2020), (iii) the number of points spent walking (i.e., when an animal is walking without feeding; Alberts et al. 2020), and (iv) the number of feeding points spent on each food type. In the “Statistical Analysis” section below we describe how these data were used to estimate the proportion of time females spent on different foraging-related activities and on different key food types in statistical models, with the goal of estimating how the odds of females engaging in these activities changed as a function of the variables described above.

## STATISTICAL ANALYSIS

### TIME SPENT ON FORAGING-RELATED ACTIVITIES

To test how group size, network density, and climate variables relate to time spent on individual foraging-related activities, we built three Generalized Estimating Equations (GEEs) using the geepack package in R (Højsgaard et al. 2006). GEEs allow for robust estimation in the presence of temporal autocorrelation, which is expected in longitudinal data where foraging behavior may be correlated across time within groups. Each of our models had one of three different binomially distributed foraging outcomes estimated from counts of focal points spent doing a behavior (successes) versus not doing a behavior (fails), which functionally represented: i) the proportion of time that all adult female baboons in a given group and year were observed to spend feeding, ii) the proportion of time adult females were observed walking without feeding, and iii) the proportion of time adult females were observed feeding relative to all foraging time (where foraging time is time spent feeding plus walking; Markham et al. 2015). We interpret the third measure to reflect the proportion of total foraging effort that female spent actually consuming food. While this metric is not a direct measure of the amount of energy gained per unit of effort, the common definition of foraging efficiency (Stacey 1986), we reasoned that, in a resource-limited system, optimizing energy balance requires minimizing time spent walking and maximizing time spent feeding (Altmann 1998; Markham et al. 2015). Baboons should therefore aim to reduce movement and increase feeding time. In our GEEs, the outcomes of proportion of time feeding, walking, and feeding per foraging time were weighted by the total number of point samples collected on adult females in a given group-year to reflect a higher certainty associated with estimates derived from a greater number of total point samples.

Each GEE included the following fixed effects: (1) a linear effect of group size in the given group-year; (2) a quadratic effect of group size (because intermediate-sized groups in this population have been shown to have smaller home ranges and shorter daily travel distances which may relate to foraging behavior; Markham et al. 2015); (3) network density in the group-year; (4) average daily rainfall during the year; (5) average maximum daily temperature during the year; and (6) an interaction between average daily rainfall and average maximum temperature. We included this interaction effect because rainfall, temperature, and food availability exhibit a very dynamic relationship in this population, which may not be fully captured by the individual effects of rainfall and temperature (see Gesquiere et al. 2024). We also included (7) a measure of “observer effort” calculated as the number of focal samples on adult females (adult males are not subjects of focal sampling) per adult individual in the group-year because we expected the intensity of observation on groups to be correlated with network density, a predictor of interest in our models. By including observer effort as an additional fixed effect our goal was to prevent any variation in the response variable explained by observer effort from being mistakenly attributed to network density, effectively controlling the estimated effects of network density for variation in the per capita intensity with which social interactions are sampled.

In each GEE, social group identity was entered as a repeated measure, and data within groups were ordered by hydrological year. We used a first order autoregressive structure to account for temporal autocorrelation among measurements (Brockwell & Davis 1991; Box et al. 2015). This correlation structure resulted in the lowest QIC score of all tested correlation structures in our models. In all models, the range of values for fixed effect variables were standardized by subtracting the mean and dividing by the standard deviation to ensure model convergence. Outliers were removed in all final reported models, although their inclusion did not qualitatively impact results.

### DIET COMPOSITION

In addition to testing how group size, network density, and climate variables covaried with time spent on foraging-related activities, we also tested how the physical and social environments covaried with diet composition, given that the types of foods an animal consumes will contribute to their overall energy intake. To do so, we built two GEEs using the geepack package in R (Højsgaard et al. 2006) to ask about the relationship between group social traits, environmental variables, and two different binomially distributed outcomes related to diet quality which represented: i) the proportion of time spent on grass corms, which typically require a large investment of time to harvest, and ii) the proportion of time spent on high energy foods, i.e., seeds, tree gum, or fruits. Baboons are well-known to rely on corms (underground storage organs) of grasses and sedges, which are nutritionally valuable but require considerable effort to harvest (Post 1982; Johnson 1989; Byrne et al. 1993, Alberts et al. 2005; Barrett & Henzi 2008). Furthermore, extensive feeding on corms is associated with relatively poor energetic condition in this population (Gesquiere et al. 2024), a pattern generally consistent with the common designation of corms as a low-yield, fallback food (Altmann 2009). The high energy foods — seeds, gums, and fruits, which are high in protein, fats, fibers, carbohydrates, minerals, and water — are considered a primary source of energy for baboons in Amboseli (Altmann 1998).

Each GEE included the same fixed effects as the models for foraging-related activities above: (1) a linear effect of group size in the given group-year; (2) a quadratic effect of group size in the group-year; (3) network density in the group-year; (4) average daily rainfall in the year; (5) average maximum daily temperature in the year; (6) an interaction effect between average daily rainfall and average maximum temperature; and (7) observer effort in the group-year. Data were again grouped by group identity and ordered by hydrological year. A first order autoregressive correlation structure was used to control for temporal autocorrelation after comparing QIC scores for models with different correlation structures. In all models, the range of values for fixed effect variables were standardized by subtracting the mean and dividing by the standard deviation to ensure model convergence. Outliers were removed in the final reported models, although their inclusion did not qualitatively impact results. Non-significant interactions were dropped from the final models.

### ASSESSING DIRECTION OF CAUSALITY

To evaluate potential causal pathways linking network density to foraging behavior, as depicted in Figure 1, we employed path analysis using the ‘lavaan’ package in R (Rosseel, 2012). We compared the performance of six different causal models explaining the relationship between network density and foraging variables (Figure 1; two models are depicted in each of the three panels). We constructed these models based on the results of our GEEs, accounting for relationships with other covariates (rainfall, maximum temperature, and group size). We evaluated the performance of each model using three performance metrics: Akaike Information Criterion (AIC), Comparative Fit Index (CFI), and Standardized Root Mean Square Residual (SRMR).

## RESULTS

### TIME SPENT ON FORAGING-RELATED ACTIVITIES AND DIET COMPOSITION ANALYSES

Figure 3 provides a schematic showing the relationship between predictors and all five outcomes in our GEE analyses.

**Figure 3:**
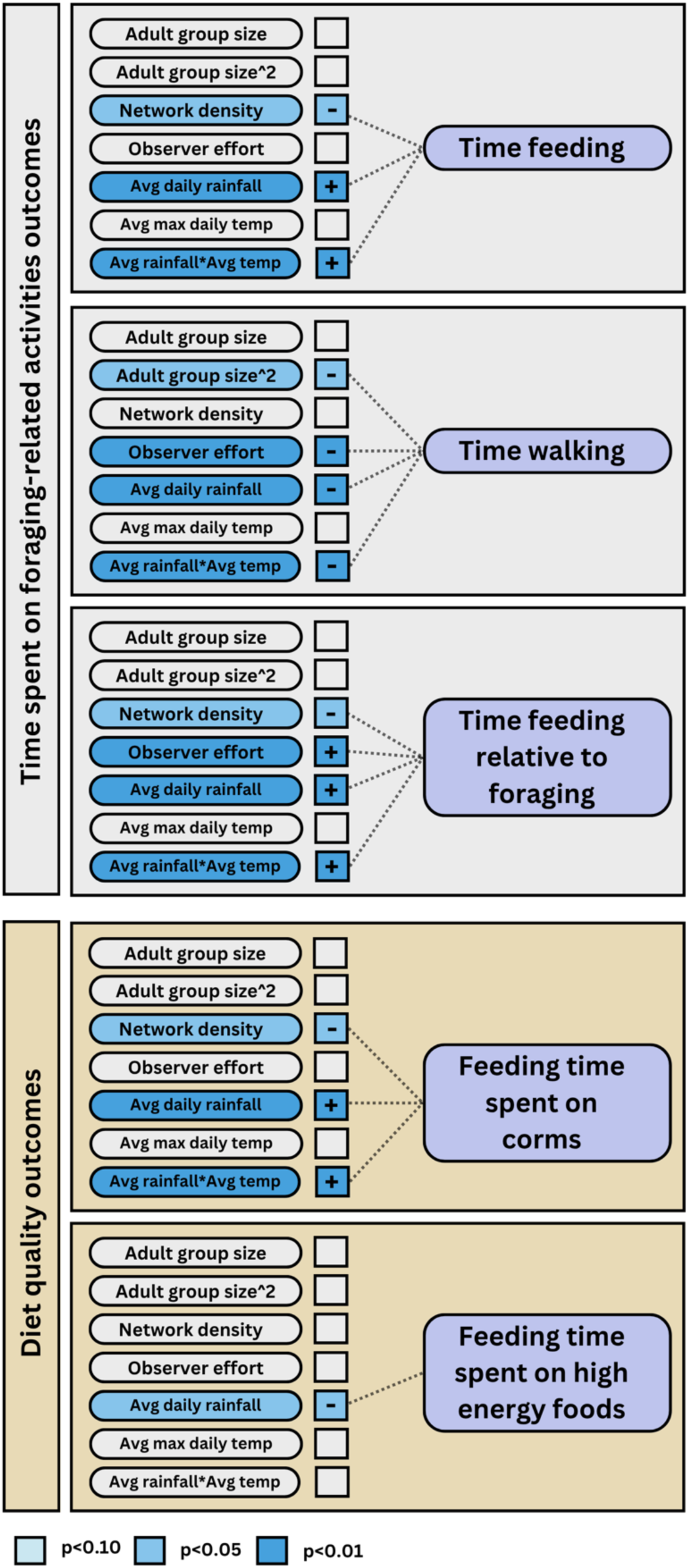
Summary of results from GEEs testing predictors of the proportion of time spent on foraging-related activities and diet composition where +/- shows the direction of the model estimate. Full results are reported in the text, in Figures 4 and 5, and in the Supplementary Material.

#### Environmental variables

Our models revealed statistically significant interactions between average daily rainfall and average maximum daily temperature that predicted time spent on foraging-related activities. During the hottest years when rainfall was high, the baboons spent more time feeding, less time walking, and more time feeding relative to walking than when rainfall was low (Figures 4A to 4C). By contrast, there was little effect of rainfall on time budgets during moderate and cooler years. For instance, for each one standard deviation increase in maximum temperature, the effect of rainfall on the odds of feeding increased by 3.6%, such that during hotter years, time spent feeding increased more dramatically with rainfall compared to cooler years (β=0.035, p=<0.001, Table S1, Figure 4A). Meanwhile, for each one standard deviation increase in maximum temperature, the effect of rainfall on the odds of walking decreased by 5%, such that during hotter years, time spent walking decreased more dramatically with rainfall compared to cooler years (β=-0.051, p=<0.001, Table S2, Figure 4B). Finally, for each one standard deviation increase in maximum temperature, the effect of rainfall on the odds of feeding during foraging increased by 5%, such that during hotter years, time spent feeding relative to all foraging time increased more dramatically with rainfall compared to cooler years (β=0.049, p=<0.001, Table S3, Figure 4C).

**Figure 4:**
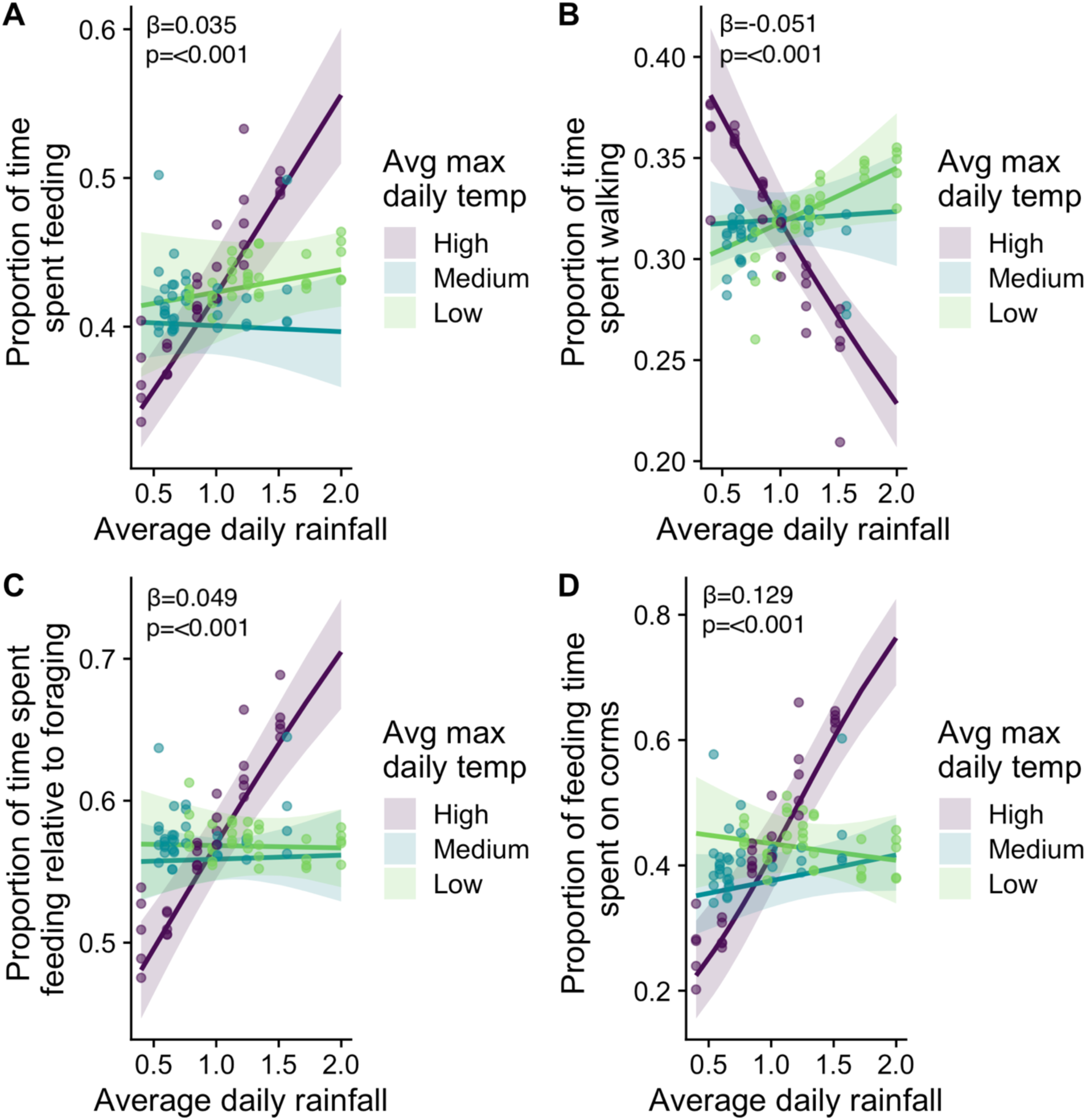
A), B), C), and D) show that rainfall and temperature interact to predict foraging outcomes. A) During hotter years, time spent feeding increases more steeply with rainfall compared to cooler years. B) During hotter years, time spent walking decreases more steeply with rainfall compared to cooler years. C) During hotter years, time spent feeding relative to all foraging time increases more steeply with rainfall compared to cooler years. D) During hotter years, the percentage of feeding time spent on corms increases more steeply with rainfall compared cooler years. Plots show the model-predicted values from GEEs modeling the relationship between average daily rainfall, average maximum daily temperature, and foraging outcomes for adult female baboons based on 100 group-years after controlling for other covariates (model results reported in Tables S1, S2, S3, and S4). Maximum temperature (a continuous variable in our GEEs) was binned into categorical tertiles for the purposes of visualizing interaction effects.

Notably, time spent feeding and diet composition covary in our dataset: during years in which groups spent more time feeding they also spent more time feeding on corms (r=0.466; Figure S4), an important fallback food that is time-consuming to process. Not surprisingly therefore, average daily rainfall and average maximum daily temperature not only interacted to predict the amount of time spent feeding overall and the amount of time spent feeding relative to foraging, but also the amount of feeding time spent eating corms. For each one standard deviation increase in maximum temperature, the effect of rainfall on the odds of eating corms increased by 13.8%. Thus, during hotter years, the percentage of feeding time spent on corms dramatically increased with rainfall compared to cooler years (β=0.129, p=<0.001, Table S4, Figure 4D). On the other hand, feeding time spent on high-energy foods was not predicted by a temperature-rainfall interaction, and instead showed a simple linear relationship with rainfall: for each one standard deviation increase in rainfall, the odds of spending time on high-energy foods decreased by approximately 7.6% (β=-0.079, p=0.022, Table S5, Figure S5).

#### Group size

Group size was significantly associated with only one of the foraging outcomes we measured: females in intermediate-sized groups spent more time walking than females in larger and smaller groups. The odds of walking (versus not walking) changed by approximately 3.4% for each one standard deviation increase in adult group size (equivalent to approximately nine additional adults), peaking in intermediate-sized groups of approximately 25 adults (β=-0.035, p=0.042; Table S2, Figure 5A). Group size was not a significant predictor of other foraging outcomes (Tables S1, S3, S4, and S5).

**Figure 5:**
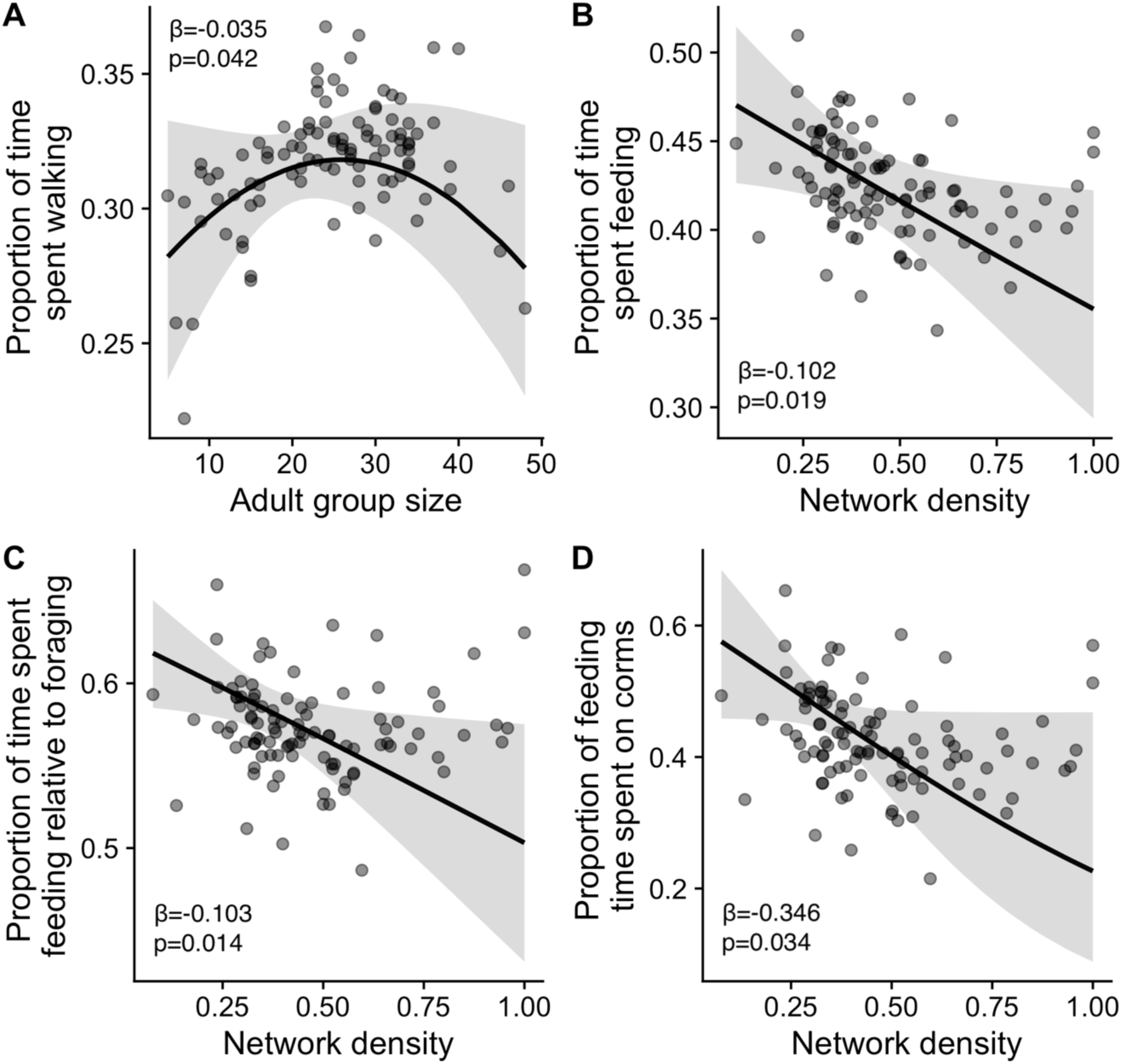
A) Intermediate-sized groups were associated with more time walking (model results reported in Table S2). B) Denser grooming networks were associated with less time feeding (model results reported in Table S1). C) Denser grooming networks were associated with less time feeding relative to foraging (model results reported in Table S3). D) Denser grooming networks were associated with less feeding time spent on fallback foods (i.e., corms) (model results reported in Table S4). Each panel shows the model-predicted trendline from a GEE modeling the relationship between a social measure and the response variable of interest, based on 100 group-years after controlling for other covariates.

#### Network density

Females in dense social networks spent considerably less proportional time feeding overall than those in sparse social networks, controlling for group size. Specifically, for each unit increase in the standardized measure of network density, the odds of feeding decreased by about 9.7% (β=-0.102, p=0.019, Table S1, Figure 5B). Female in dense networks also spent less proportional time feeding relative to foraging than those in sparse social networks: for each unit increase in standardized network density, the odds of feeding while foraging decreased by about 9.8% (β=-0.103, p=0.014, Table S3, Figure 5C). Given that network density was not significantly associated with the proportion of time spent walking (Table S2), this decrease in feeding relative to foraging time (where foraging is time spent feeding plus walking) in denser networks is likely driven by the decreased time spent feeding in these groups, as opposed to increases in time spent walking without feeding.

Females in dense social networks also relied substantially less on grass corms than those in sparse social networks, suggesting that they may have had a foraging advantage compared to females in groups with sparser networks (since corms take time and energy to process). Specifically, for each one standard deviation increase in network density, the odds of eating corms decreased by 29.2% (β=-0.346, p=0.034, Table S4, Figure 5D). Network density was not associated with the proportion of feeding time spent on high energy foods (Tables S5).

#### Observer effort

We included observer effort as a covariate in all models to control for its effect on network density, one of our primary variables of interest. This measure had an unexpected relationship with several foraging outcomes, which we show and interpret in the Supplementary Materials.

### ASSESSING DIRECTION OF CAUSALITY

The causal relationships between network density and its associated foraging outcomes could take several, non-mutually exclusive, forms, a subset of which are shown in Figure 1. One possible scenario involves foraging trade-offs, in which diet composition (in this case the proportion of the diet that is corms) and (directly or indirectly) time spent feeding causally affect social network density (Figure 1A and 1B). Another possible scenario involves social benefits, in which network density causally affects diet composition and (directly or indirectly) time spent feeding (e.g., via information sharing and success in inter-group competition; Wright et al. 2001; Silk 2007; Burkart 2017) (Figure 1C and 1D). Furthermore, both processes may act simultaneously, in which case causal arrows flow from network density to foraging outcomes and from foraging outcomes to network density (Figure 1E and 1F).

Our path analysis showed that the best supported model based on AIC, CFI, and SRMR was the causal path detailed in Figure 1E, where the proportion of corms being consumed affects network density (the foraging trade-off hypothesis) and network density separately affects time spent feeding (the social benefits hypothesis), with an AIC of 1820, CFI of 0.963, and SRMR of 0.041 (Figure 6, Tables S6 and S7). That is, our analysis supports the idea that both foraging trade-offs and social benefits may be acting in this study population. However the model in Figure 1A, was within two ΔAIC of the top model (Table S6). This second-best model, which is statistically indistinguishable from the top model based on its AIC, posits that the proportion of corms being consumed has a causal effect on network density (foraging trade-off hypothesis), and that the relationship between network density and time spent feeding is indirect. Both models therefore support a direct effect of diet composition (i.e., proportion of feeding time spent on corms) on network density, providing support for the foraging trade-off hypothesis. Importantly, unlike our GEE models these causal models did not accommodate information on group identity and time series, limiting the comparability and interpretability of these models. The limited number of individual group-years for each group prevented us from building more complex causal models that included this information.

**Figure 6:**
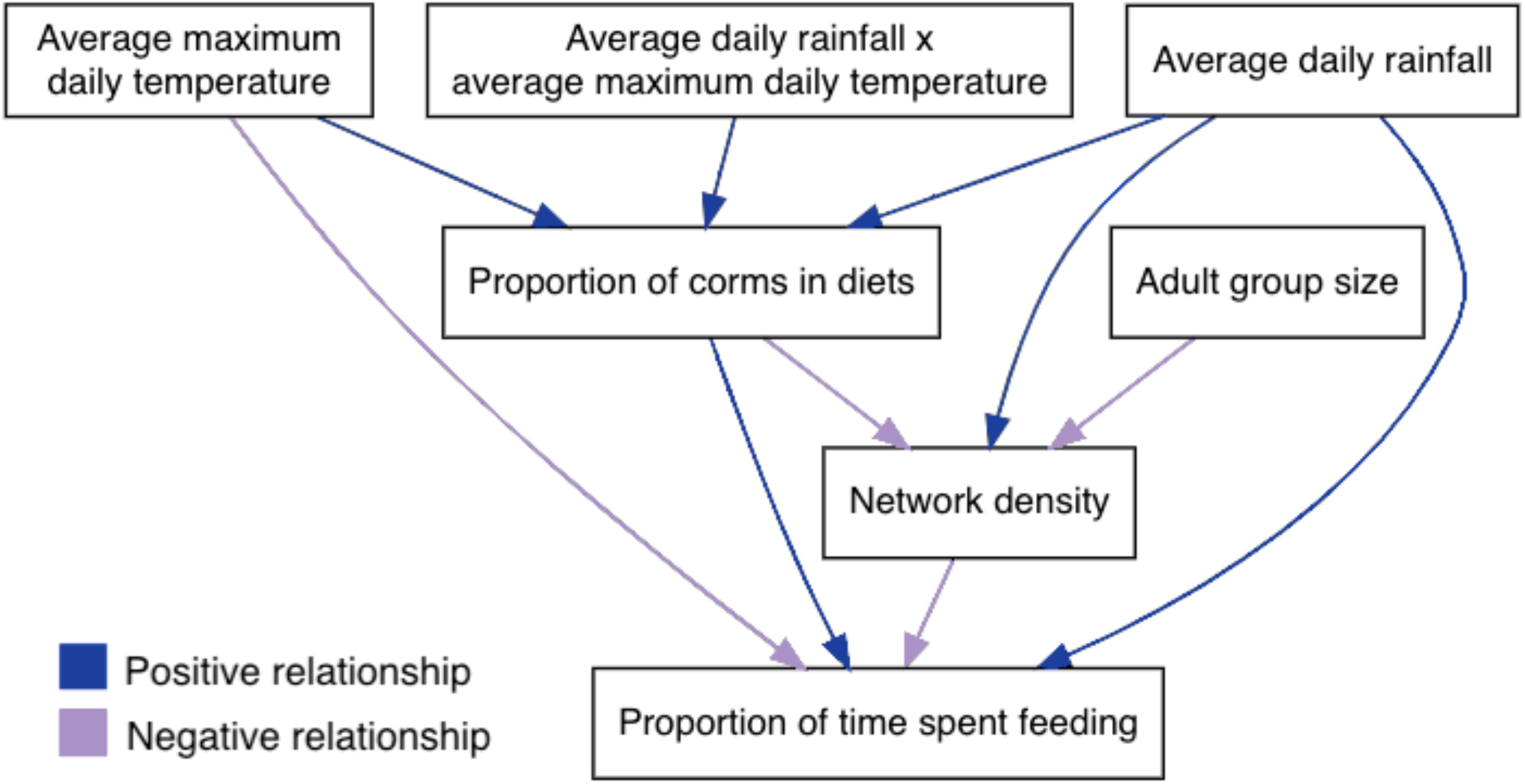
Top performing model in path analysis with effect sizes AIC=1820, CFI: 0.963, and SRMR: 0.041 (see Tables S6 and S7). This model supports a causal effect of diet composition (i.e., the proportion of feeding time spent on fallback foods) on network density, and also suggests that increased network density separately drives a reduction in time spent feeding.

## DISCUSSION

Our analyses showed that conditions in both the physical and social environment were associated with the amount of time animals spent on foraging-related activities and with diet composition. Conditions that were associated with investing more in directly obtaining food (i.e., more time spent feeding and less time spent walking without feeding) also tended to be associated with a high proportion of corms in the diet. Corms are a common fallback food, suggesting that diet composition likely plays a large role in determining how animals allocate their time-budgets in this populaton, e.g., foods that are time consuming to eat lead animals to devote more of their time budget to feeding. For instance, during hot and rainy years animals spent a high proportion of feeding time on corms, and also spent more time spent feeding, more time feeding relative to foraging, and less time walking. These results indicate that when feeding on corms, which are considered to be nutritionally low-yield, animals must spend more time feeding to meet their nutritional demands. However, in these years feeding more on corms also means that animals spend less time and energy searching for food (i.e., walking).

While baboons spent the most time feeding on corms in hot and rainy years, they spent the least time feeding on corms in hot and dry years, even less than cool years (see the steep slope for high temperature years in Figure 4D). This pattern during hot years, in which fewer corms are consumed during years with less rain, is the opposite of the pattern we observe within years, where corms are most commonly eaten during the annual “long dry season” between May and October (Alberts et al. 2005; Gesquiere et al. 2024). We believe that this difference between within-year patterns of corm consumption and our between-year analysis arises from the effects of drought. In typical years, the hottest months are the rainiest months, but during drought years, the hottest months are dry, and even hotter than during typical years. Thus the hot and dry years in our dataset represent those years with severe drought conditions. Although corms are an abundant resource during short periods of dryness (i.e., dry seasons), we investigated corm condition at the end of a two-year drought in Amboseli and found that corms from areas that received no rain were completely dried out (Figure S6). Therefore, while corms are usually a reliable fallback food during short bouts of dryness, this may not be the case during drought years. This result represents a previously unknown constraint on the behavioral flexibility of baboon foraging patterns.

Socially dense grooming networks were associated with less time spent feeding than socially sparse networks and individuals in these sparse networks also spent more feeding time on corms. Notably, these observed relationships between network density and foraging outcomes have multiple potential explanations that are difficult to disentangle. We tested two hypotheses explaining the relationship between network density and foraging: i) the social benefits hypothesis, which predicts that a cohesive group structure may improve the efficiency of group feeding and give individuals in the group access to better resources and ii) the foraging trade-off hypothesis which predicts individuals may create denser social networks when they experience better environmental conditions that enable them to forage more efficiently and eat fewer low quality foods. A path analysis exploring these hypotheses revealed two top models that support a causal effect of the proportion of feeding time spent on corms (a fallback food) on network density. Thus, both top models support the foraging trade-off hypothesis, suggesting that females in groups with higher quality diets (less feeding time spent on corms) have more time and energy to create social connections. Nonetheless, we also found some support for the social benefits hypothesis: one of our top models suggested that increased network density may separately drive a reduction in time spent feeding. Notably, our models were limited in their potential for causal inference given that we could not test all possible external predictors of network density and foraging; unmeasured third variables could also mediate the observed effects. Moreover, due to sample size limitations we could not incorporate information about group identity and temporal autocorrelation into the causal models. Finally, our simple path models assumed unidirectional causality, an assumption that could be violated if feedbacks between network density and particular foraging variables exist. Thus, conclusions from these models should be interpreted as tentative and more robust models should be tested when enough data are available.

A causal scenario in which an increased use of fallback foods leads to decreased network density is perhaps the more parsimonious of the two possible explanations for a relationship between diet quality and network density. This is because the alternative scenario, in which individuals accrue foraging benefits from social relationships, would require more complex cognitive and social processes than one in which individuals simply reduce socializing when they must feed more or are feeding on more spatially distributed foods. Notably, a higher network density means that more potential social links in a group are realized. Thus, if the foraging trade-off hypothesis explains these results, it would mean individuals with better diets not only spend more time and energy socializing, but they also expand their social networks, socializing with a broader range of individuals, as opposed to only investing additional time into their existing bonds.

Future research in behavioral physiology could lend additional insight into the mechanisms linking network density and foraging outcomes. For instance, analyses of thyroid hormones, which are a sensitive measure of energy expended relative to energy acquired (Flier et al. 2000; Markham & Gesquiere 2017), combined with intensive behavioral sampling, would enable short-term trends in social behavior to be compared against short-term trends in energetic levels to evaluate if one systematically preceeds the other. Decreases in foraging time and improved energetic condition that precede, even by a few days, changes in social network density would provide support for the foraging tradeoffs hypothesis. In contrast, increases in social network density that precede decreases in time spent feeding and improvements in energetic states would favor the social benefits hypothesis. Such cross-lagged analyses would require exceptionally concentrated behavioral data to ask about delayed effects on the scale of e.g., days, but represent an alternative method of causal inference to path analysis.

Markham et al. (2015) showed that intermediate-sized groups in this population have smaller home range size, shorter daily distance traveled, and more even patterns of space use than large and small groups, but that large groups spend more time foraging (feeding plus walking) than intermediate and small groups. Here, we found no linear relationships between group size and foraging-related behaviors (Tables S1 to S5), but intermediate-sized groups in our study actually spent more time walking without feeding (Table S2), suggesting that they expended more energy searching for food than the smallest and largest groups. In other words, while previous studies detected a benefit of intermediate-sized groups in terms of space use, we found these groups may expend more energy foraging in the spaces they occupy. There is some indication, although not statistically significant in our analysis, that intermediate-sized groups rely less on corms as a food resource (Table S4), which might mean that the additional time they spent seeking food is directed towards higher-yield foods that are less abundant.

In conclusion, we found that conditions in both the social and physical environment are associated with the foraging outcomes achieved by adult female baboons. Conditions that favor focus on low-yield fallback foods result in individuals spending more time feeding to meet their nutritional requirements. However, eating these lower quality foods may also allow animals to reduce movement under some conditions (e.g., when it is hot and rainy). While predictors like temperature and rainfall have clear causal effects on foraging outcomes, network density may affect or be affected by foraging outcomes. If diet composition affects network density (as suggested in our path analysis), the increased time spent feeding when the diet includes a high proportion of corms may take away from time and energy left to invest in maintaining social connections, thus leading to lower network densities in these groups. If true, future work could investigate how increasing variable climate conditions affect food availability and, in turn, social networks in wild animal populations.

## DATA AVAILABILITY STATEMENT

Data and code will be made available upon publication.

## ACKNOWLEDGMENTS

We gratefully acknowledge the support of the National Science Foundation and the National Institutes of Health for the majority of the data represented here, through multiple grants over the years, including currently active NIH grants R01AG53330, R01AG053308, R01AG071684, R01AG075914, and R61AG078470. Current support for field-based data collection also comes from the Max Planck Institute for Evolutionary Anthropology, and we thank Duke University, Princeton University, and the University of Notre Dame for financial and logistical support. The research activity for this specific project was additionally supported by the Natural Sciences and Engineering Research Council of Canada (NSERC) and the Triangle Center for Evolutionary Medicine (TriCEM). We thank N.Z. Kerr and A.C. Markham for helpful discussions on statistical modeling. In Kenya, our research was approved by the Wildlife Research Training Institute (WRTI), Kenya Wildlife Service (KWS), the National Commission for Science, Technology, and Innovation (NACOSTI), and the National Environment Management Authority (NEMA). We also thank the University of Nairobi, the Kenya Institute of Primate Research (KIPRE), the National Museums of Kenya, the members of the Amboseli-Longido pastoralist communities, the Enduimet Wildlife Management Area, Ker & Downey Safaris, Air Kenya, and Safarilink for their cooperation and assistance in the field. Particular thanks go to the Amboseli Baboon Project long-term field team (R.S. Mututua, S. Sayialel, I.L. Siodi, and L. Musembei) for data collection, and to T. Wango and V. Oudu for their assistance in Nairobi. The baboon project database, Babase, has been expertly managed by N. Learn, J. Gordon, and W. Wilbur. Database design and programming are provided by K. Pinc. This research was approved by the IACUC at Duke University, University of Notre Dame, and Princeton University and the Ethics Council of the Max Planck Society and adhered to all the laws and guidelines of Kenya. For a complete set of acknowledgments of funding sources, logistical assistance, and data collection and management, please visit http://amboselibaboons.nd.edu/acknowledgements/.

