## Supplementary Materials for "Group social conditions and environment predict foraging behavior in wild baboons"

### CORRELATIONS BETWEEN OBSERVER EFFORT AND FORAGING OUTCOMES

Observer effort was included as a covariate in generalized estimating equations (GEEs) to control for its effect on observed network density. This effect arises because the team of observers for the Amboseli Baboon Research Project stays relatively constant in size while baboon groups vary in size. Data on social interactions are collected as each observer moves systematically through the group while carrying out 10-minute focal follows according to a predefined, randomized list of focal animals. While the observers record all grooming interactions in their line of sight, whether or not they involve the focal animal, any given social interaction is inevitably less likely to be observed in a large group than in a small group. By including a measure of observer effort in our models we control for variance in network density that is explained by variance in the per capita sampling intensity. We measured observer effort as the number of 10-minute focal samples taken on adult females per adult individual in a group.

We found that observer effort was associated with the outcomes of time spent walking (Table S2) and time spent feeding relative to total time foraging (Table S3). After some investigation we believe the most parsimonious explanation for this relationship stems from the fact that poor sampling conditions, which lead to observers losing sight of focal animals and halting focal sampling (resulting in lower observer effort) are more likely to occur when animals are engaged in certain activities like walking. For example, when animals are walking, focal individuals can be more difficult to locate, and observers are more likely to lose sight of either their focal animal or the entire group (and thus stop sampling all together). Therefore, we believe that when animals are walking, observers are more likely to have a focal individual out of sight, complete less focals altogether, and observe fewer ongoing social interactions.

### SUPPLEMENTARY TABLES

Table S1: Predictors of proportion of total time adult female baboons spend feeding each year. Results from a GEE model across 100 group years between 2000 and 2021. Data grouped by group identity and ordered by hydrological year.

| term | estimate | std error | wald | p value |
|---|---|---|---|---|
| **(Intercept)** | -0.306 | 0.042 | 54.360 | <0.001** |
| **Adult group size** | -0.036 | 0.058 | 0.380 | 0.537 |
| **Adult group size^2** | 0.020 | 0.022 | 0.760 | 0.383 |
| **Network density** | -0.102 | 0.044 | 5.490 | 0.019* |
| **Observer effort** | 0.017 | 0.024 | 0.520 | 0.469 |
| **Avg daily rainfall** | 0.100 | 0.016 | 39.790 | <0.001** |
| **Avg max daily temp** | -0.008 | 0.016 | 0.240 | 0.625 |
| **Avg daily rainfall x avg max daily temp** | 0.035 | 0.007 | 23.680 | <0.001** |

Note: continuous variables were scaled by subtracting the mean and dividing by the standard deviation. +p<0.1; * p<0.05; ** p<0.01.

Table S2: Predictors of the proportion of total time adult female baboons spend walking each year. Results from a GEE model across 100 group years between 2000 and 2021. Data grouped by group identity and ordered by hydrological year.

| term | estimate | std error | wald | p value |
|---|---|---|---|---|
| **(Intercept)** | -0.759 | 0.034 | 490.510 | <0.001** |
| **Adult group size** | 0.008 | 0.047 | 0.030 | 0.866 |
| **Adult group size^2** | -0.035 | 0.017 | 4.150 | 0.042* |
| **Network density** | 0.045 | 0.036 | 1.590 | 0.208 |
| **Observer effort** | -0.074 | 0.023 | 10.710 | 0.001** |
| **Avg daily rainfall** | -0.048 | 0.012 | 15.790 | <0.001** |
| **Avg max daily tem** | 0.008 | 0.011 | 0.500 | 0.479 |
| **Avg daily rainfall x avg max daily temp** | -0.051 | 0.011 | 20.560 | <0.001** |

Note: continuous variables were scaled by subtracting the mean and dividing by the standard deviation. +p<0.1; * p<0.05; ** p<0.01.

Table S3: Predictors of the proportion of time adult female baboons spend feeding relative to total time foraging each year (defined as feeding plus walking). Results from a GEE model using 100 group years between 2000 and 2021. Data grouped by group identity and ordered by hydrological year.

| term | estimate | std error | wald | p value |
|---|---|---|---|---|
| **(Intercept)** | 0.290 | 0.037 | 62.240 | 0.001** |
| **Adult group size** | -0.041 | 0.049 | 0.680 | 0.411 |
| **Adult group size^2** | 0.033 | 0.021 | 2.510 | 0.113 |
| **Network density** | -0.103 | 0.042 | 6.010 | 0.014* |
| **Observer effort** | 0.061 | 0.019 | 10.280 | 0.001** |
| **Avg daily rainfall** | 0.090 | 0.016 | 32.500 | <0.001** |
| **Avg max daily temp** | -0.006 | 0.015 | 0.140 | 0.711 |
| **Avg daily rainfall x avg max daily temp** | 0.049 | 0.010 | 23.000 | <0.001** |

Note: continuous variables were scaled by subtracting the mean and dividing by the standard deviation. +p<0.1; * p<0.05; ** p<0.01.

Table S4: Predictors of the proportion of time adult female baboons spend feeding on fallback foods (i.e., corms) relative to total time feeding each year. Results from a GEE model using 100 group years between 2000 and 2021. Data grouped by group identity and ordered by hydrological year.

| term | estimate | std error | wald | p value |
|---|---|---|---|---|
| **(Intercept)** | -0.355 | 0.112 | 10.000 | 0.002** |
| **Adult group size** | -0.172 | 0.169 | 1.040 | 0.307 |
| **Adult group size^2** | 0.096 | 0.059 | 2.670 | 0.102 |
| **Network density** | -0.346 | 0.163 | 4.520 | 0.034* |
| **Observer effort** | 0.084 | 0.060 | 1.970 | 0.161 |
| **Avg daily rainfall** | 0.263 | 0.044 | 35.900 | <0.001** |
| **Avg max daily temp** | -0.038 | 0.055 | 0.470 | 0.493 |
| **Avg daily rainfall x avg max daily temp** | 0.129 | 0.034 | 14.250 | <0.001** |

Note: continuous variables were scaled by subtracting the mean and dividing by the standard deviation. +p<0.1; * p<0.05; ** p<0.01.

Table S5: Predictors of the proportion of time adult female baboons spend feeding on high energy foods relative to total time feeding each year. Results from a GEE model using 100 group years between 2000 and 2021. Data grouped by group identity and ordered by hydrological year. See text for definition of high energy foods.

| term | estimate | std error | wald | p value |
|---|---|---|---|---|
| **(Intercept)** | -0.991 | 0.073 | 184.180 | <0.001** |
| **Adult group size** | 0.085 | 0.154 | 0.310 | 0.579 |
| **Adult group size^2** | -0.042 | 0.054 | 0.620 | 0.431 |
| **Network density** | 0.142 | 0.122 | 1.340 | 0.246 |
| **Observer effort** | -0.045 | 0.074 | 0.370 | 0.544 |
| **Avg daily rainfall** | -0.079 | 0.035 | 5.260 | 0.022* |
| **Avg max daily temp** | -0.014 | 0.034 | 0.160 | 0.689 |

Note: continuous variables were scaled by subtracting the mean and dividing by the standard deviation. +p<0.1; * p<0.05; ** p<0.01.

Table S6: Comparison of performance metrics for six causal models depicted in Figure 1.

| Model | AIC | CFI | SRMR |
|---|---|---|---|
| **A** | 1822 | 0.948 | 0.045 |
| **B** | 1823 | 0.936 | 0.047 |
| **C** | 1830 | 0.862 | 0.073 |
| **D** | 1826 | 0.910 | 0.062 |
| **E** | 1820 | 0.963 | 0.041 |
| **F** | 1826 | 0.907 | 0.057 |

Table S7: Estimates from the best supported causal model among those shown in Figure 1E, describing the directional relationships among foraging variables, group size, network density (controlled for observer effort), and environmental variables. Note: The causal models used here did not incorporate group size dynamics or temporal structure, which limits direct comparability with models presented in earlier sections. As such, these estimates are subject to interpretive limitations.

| **paths** | **estimate** | **std error** | **z value** | **p value** |
|---|---|---|---|---|
| **prop diet corms ~** | | | | |
| avg rainfall | 0.269 | 0.102 | 2.640 | 0.008 |
| max temp | 0.063 | 0.105 | 0.605 | 0.545 |
| rain x max temp | 0.346 | 0.071 | 4.897 | <0.001 |
| **prop time feeding ~** | | | | |
| avg rainfall | 0.327 | 0.099 | 3.313 | 0.001 |
| max temp | -0.155 | 0.094 | -1.659 | 0.097 |
| prop diet corms | 0.396 | 0.084 | 4.727 | <0.001 |
| **network density ~** | | | | |
| avg rainfall | 0.222 | 0.049 | 4.529 | <0.001 |
| adult group size | -0.238 | 0.049 | -4.902 | <0.001 |
| prop diet corms | -0.147 | 0.049 | -2.989 | 0.003 |
| **prop time feeding ~** | | | | |
| network density | -0.383 | 0.147 | -2.597 | 0.009 |

**SUPPLEMENTARY FIGURES**

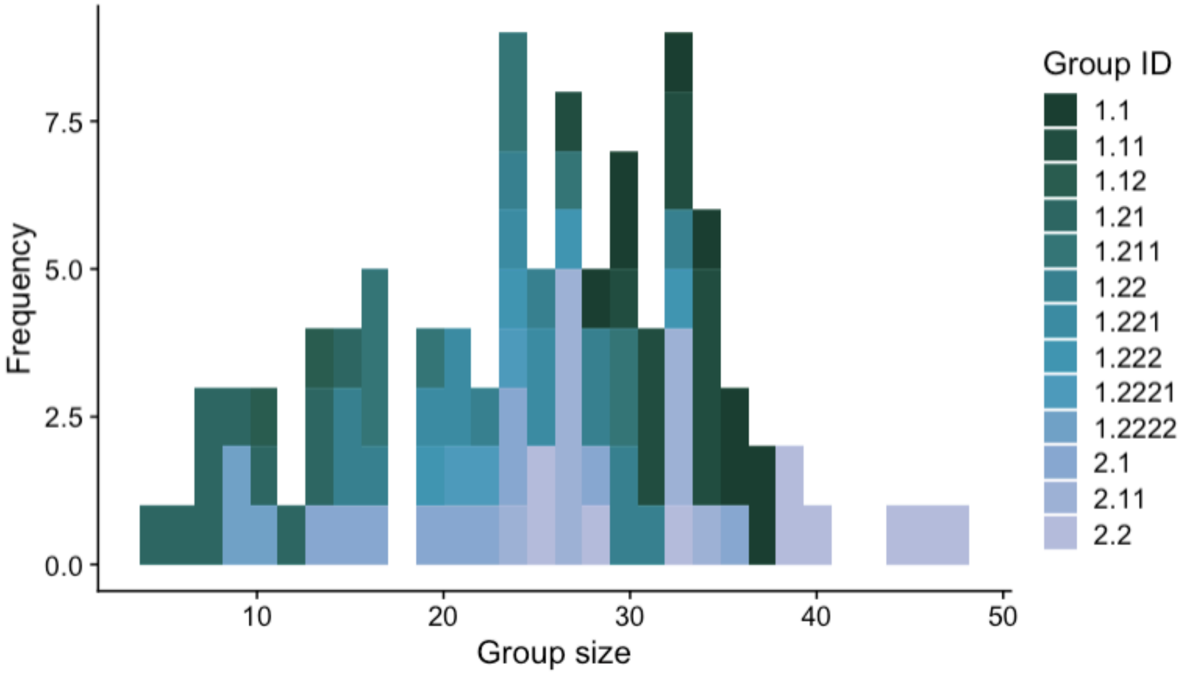


Figure S1: Histogram of group size as the total number of adults in a group in a given year for the 100 group-years included in our analyses. Color indicates group identity.

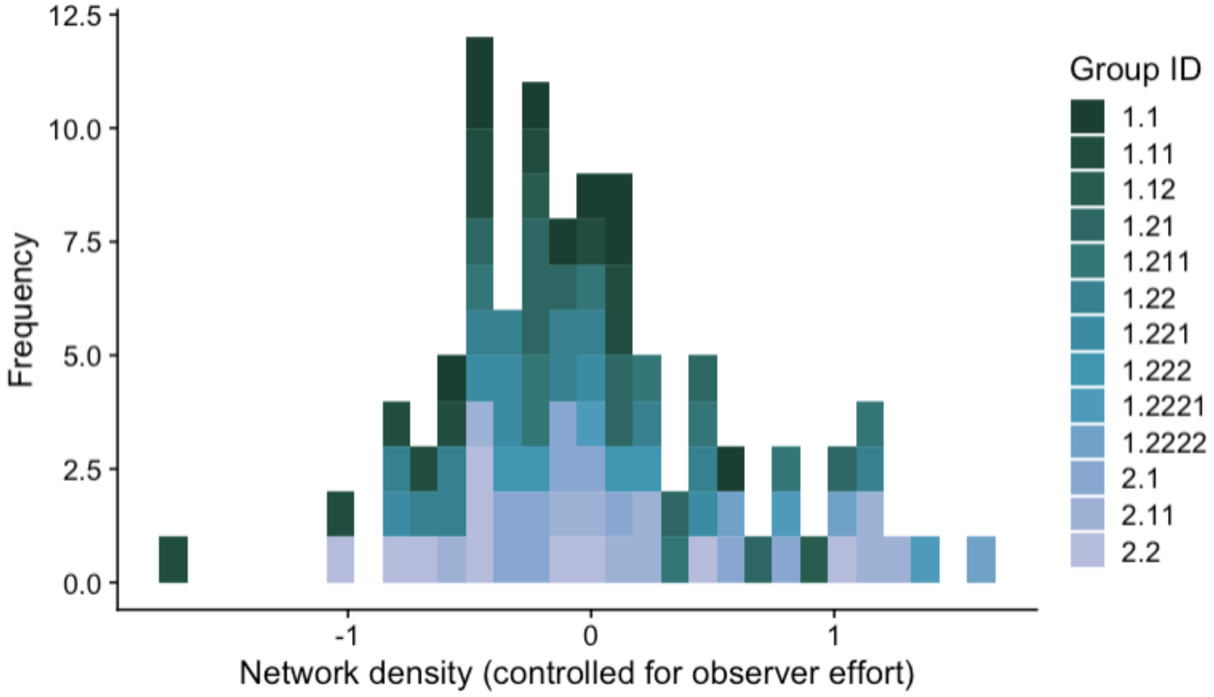


Figure S2: Histogram of annual grooming network density values for the 100 group-years of data included in our analyses. Our annual measures of network density controlled for observer effort by taking residuals from a ln-ln linear regression model of network density on the number of 10-minute focal follows conducted by observers per adult in a group. Color indicates group identity.

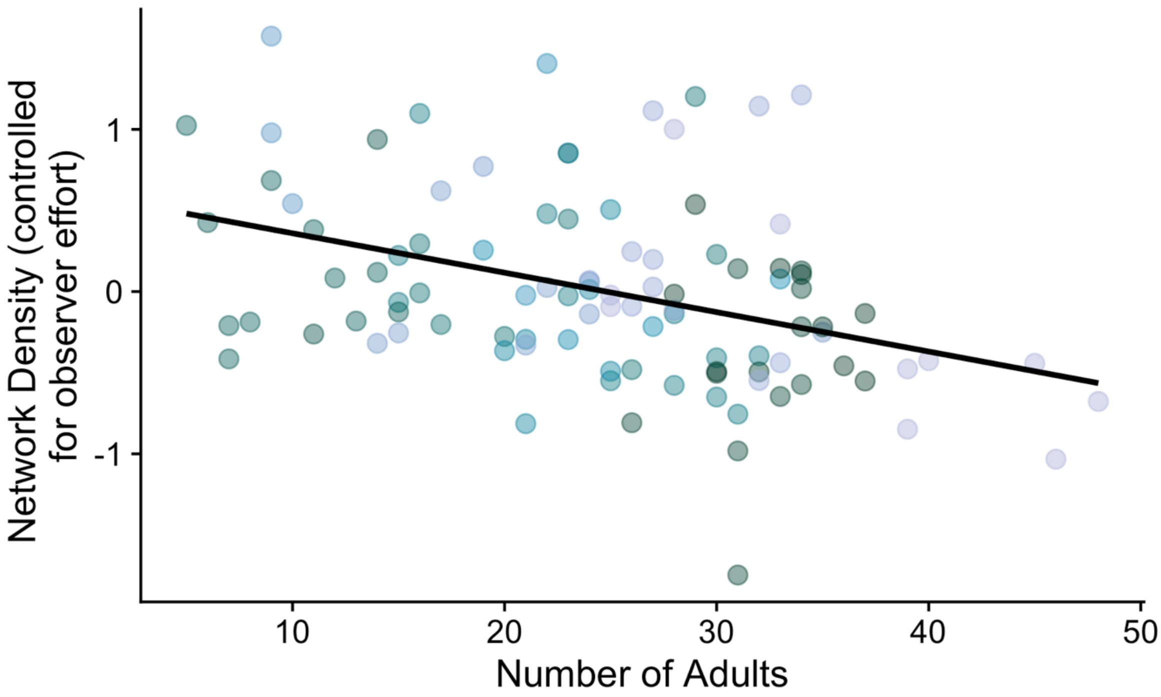


Figure S3: Network density decreases as the number of adults in a group increases (r=-0.386). X-axis shows the number of adults in a group in a given year. Y-axis shows the residuals of a linear regression of network density on observer effort. Each point represents one group-year (n=100 group years). Points of the same color come from the same group in different group years.

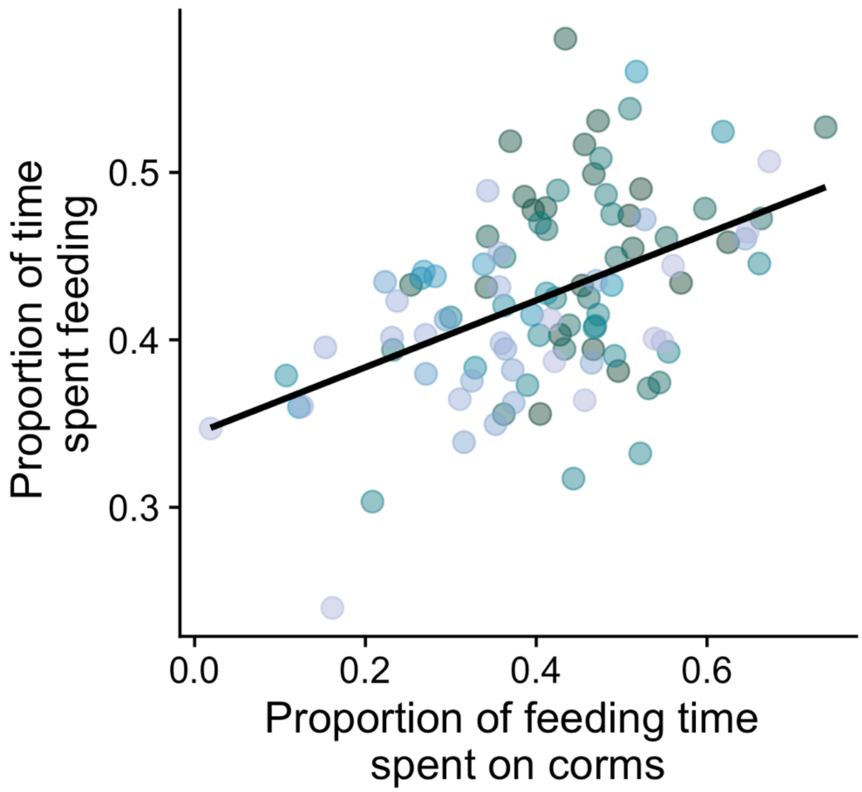


Figure S4: Time spent feeding increases with the proportion of feeding time spent on corms (r= 0.466). Points of the same color come from the same group in different group years (n=100 group years).

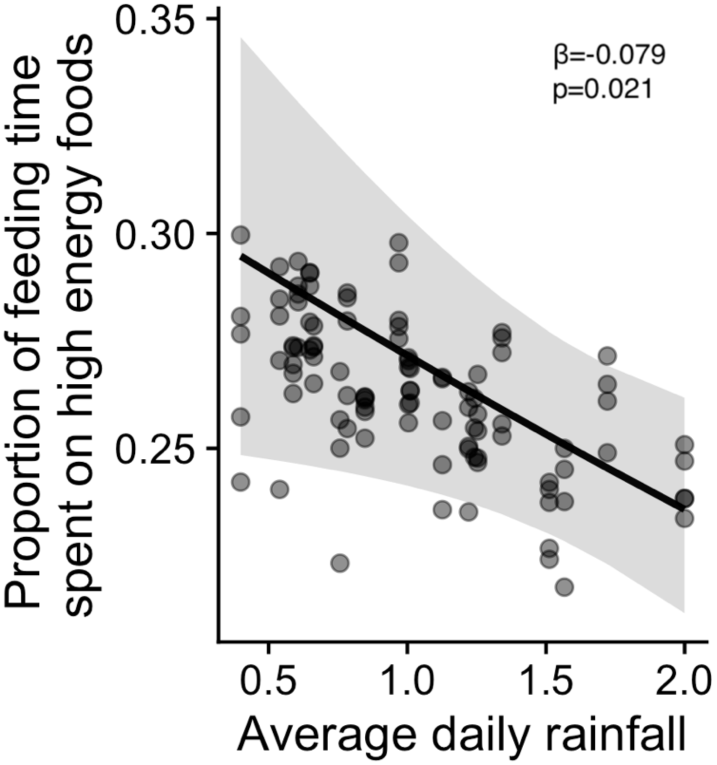


Figure S5: During years with high rainfall adult females spend less time eating high energy foods. Plot shows the model-predicted values from a GEE modeling the relationship between average daily rainfall and proportion of time spent on high energy foods for adult female baboons across 100 group years after controlling for other covariates (model results reported in Table S5).

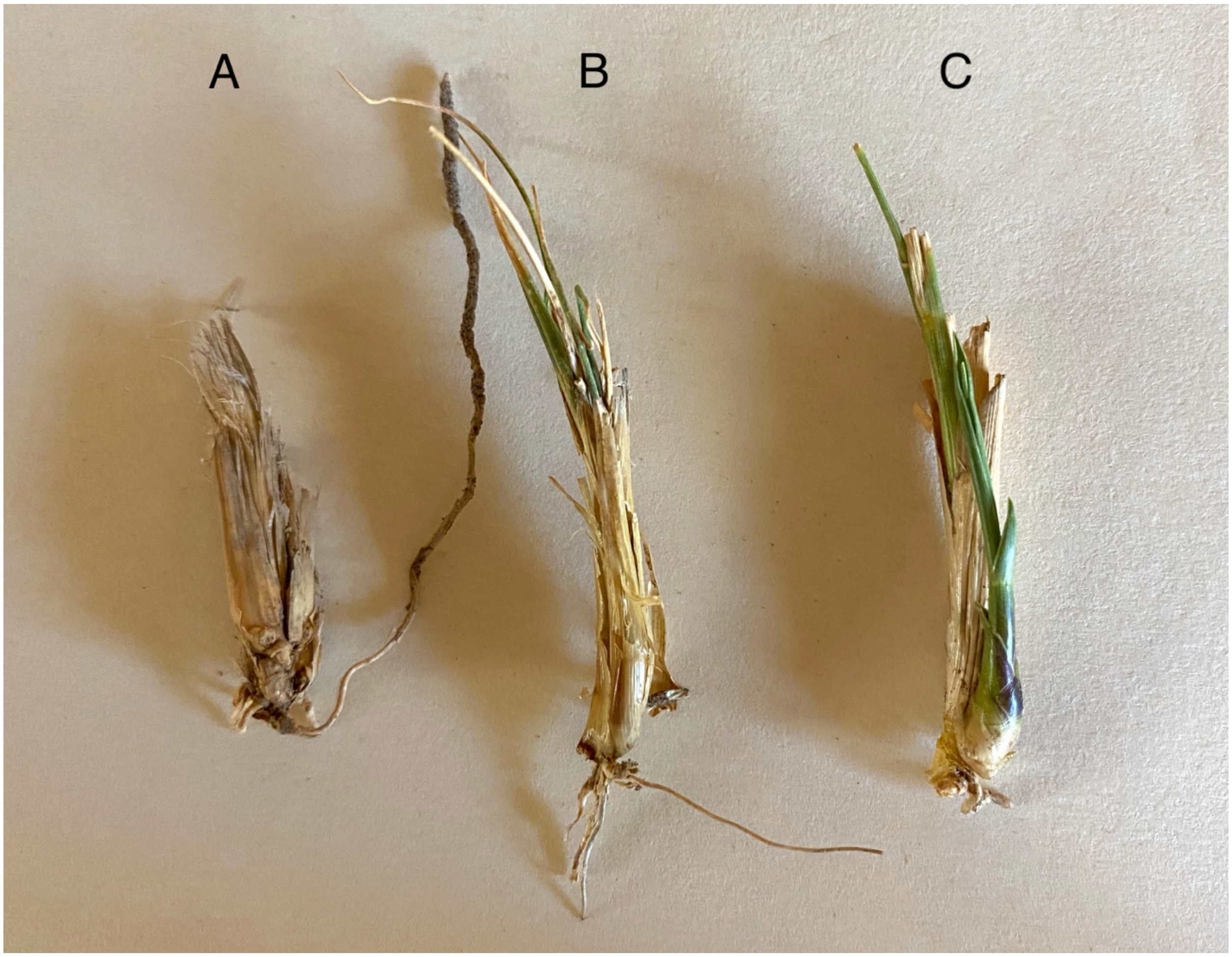


Figure S6: Corms photographed at start of the long rainy season (March) in 2023. This season marked the end of a two-year drought in Amboseli. Corms were taken from three different locations: A) an area that had not yet began receiving rain, B) an area that had received one instance of light rain, and C) and area that had received two to three instances of light rain.
